# Low-pathogenic virus induced immunity against TBEV protects mice from disease but not from virus entry into the CNS

**DOI:** 10.1101/2021.01.11.426200

**Authors:** Monique Petry, Martin Palus, Eva Leitzen, Johanna Gracia Mitterreiter, Bei Huang, Andrea Kröger, Georges M.G.M Verjans, Wolfgang Baumgärtner, Guus F. Rimmelzwaan, Daniel Růžek, Albert Osterhaus, Chittappen Kandiyil Prajeeth

**Author notes:** Correspondence to Dr. Chittappen Kandiyil Prajeeth.

## Abstract

Tick-borne encephalitis virus (TBEV) is a leading cause of vector-borne viral encephalitis with expanding endemic regions across Europe. Although currently used inactivated whole virus vaccines are effective, vaccination breakthroughs have been reported for which the reasons are unclear. In this study we tested in mice the efficacy of pre-infection with a closely related low-virulent flavivirus, Langat virus (LGTV strain TP21), or a naturally avirulent TBEV strain (TBEV-280) in providing protection against lethal infection with the highly virulent TBEV strain TBEV-Hypr (referred to as TBEV-Hypr). LGTV has been evaluated as an experimental live vaccine against TBE, but further development was abandoned due to too high residual pathogenicity of a LGTV-based vaccine. Here we show that prior infection with TP21 or TBEV-280 is efficient in protecting mice from lethal TBEV-Hypr challenge. Histopathological analysis of brains from non-immunized control mice revealed neuronal TBEV infection and necrosis. Neuroinflammation, gliosis and neuronal necrosis was however also observed in some of the TP21 and TBEV-280 pre-infected mice although at reduced frequency as compared to the non-immunized TBEV-Hypr infected control mice. Interestingly, qPCR detected the presence of viral RNA in the brains and spinal cord of both TP21 and TBEV-280 immunized mice after TBEV-Hypr challenge, but significantly reduced compared to mock-immunized mice. Our results indicate that although TBEV-Hypr infection is effectively controlled in the periphery upon immunization with low-virulent LGTV or naturally avirulent TBEV-280, it may still enter the CNS of these animals. These findings improve our understanding of potential causes for vaccine failure in individuals vaccinated with TBE vaccines.

## Introduction

Tick-borne encephalitis virus (TBEV) is an enveloped virus belonging to the family *Flaviviridae* which contains several other vector-transmitted viruses such as West Nile virus, yellow fever virus, dengue virus, Japanese encephalitis virus and Zika virus (ZIKV), which all may cause life threatening conditions in humans (1). TBEV is a major cause of concern in Europe and Asia with currently expanding endemic areas (2). TBEV is primarily transmitted by a tick-bite of Ixodes species, which can lead to strain-dependent outcome of illness (3,4). The clinical course of infections caused by TBEV strains circulating in Europe often shows a biphasic pattern. Approximately 7-14 days after infection, first flu-like symptoms may appear. After an asymptomatic phase, 20-30% of the patients develop a second phase with headache, high fever, and neurological symptoms as a consequence of severe meningitis and meningoenephalomyelitis. Of these patients, 2% exhibit long-lasting neurological sequelae (5,6). Preventive vaccination is the only available specific intervention method for TBE to date. Current TBEV vaccines are based on formalin inactivated whole virus preparations, which have proven to be effective in conferring temporary protection. Therefore repeated vaccinations are needed to induce satisfactory immunity with booster doses being required every 5-10 years (6–9). Live attenuated vaccines have been proven to confer better protection, nevertheless associated safety risks have discouraged their usage (10,11). LGTV, a naturally occurring low-virulent flavivirus, that has high amino acid sequence identity with TBEV (∼80%), has shown potential as a live attenuated vaccine candidate. In Russia LGTV was used in 1970s to vaccinate against TBE, which was regarded highly successful until among vaccinees cases displaying neurological symptoms (about 1:20.000) were observed (2,12–15). Apparently, this LGTV-based vaccine was under-attenuated. Protection against TBEV and other flaviviruses in the periphery is largely mediated by virus-specific antibodies. In contrast, virus control within the central nervous system (CNS) is dependent on the interplay between infiltrating virus specific T cell and CNS cells. It has been shown that this interaction may have both beneficial and detrimental effects (16–18). Investigations into the pathogenesis of TBE in patients revealed that granzyme B releasing T cells and microglia cells / macrophages contribute to the tissue damage after infection, resulting in neuronal death and astrogliosis (19). It was recently shown that even low dosage LGTV infection in mice may result in astrogliosis and microglia activation in the hippocampus, suggesting that infection with non-lethal doses of flaviviruses can indeed lead to histopathological changes in the brain (20).

To gain more insights in live vaccine induced protective immunity against TBE, we studied the efficacy of LGTV and naturally avirulent member of the TBEV serogroup (TBEV-280; a strain closely related to another naturally avirulent and well-characterized strain TBEV-263 (21)) in conferring protection against challenge with highly pathogenic TBEV strain in mice. Our findings show the potential of TP21 and TBEV-280 in providing protection against infection with the highly pathogenic TBEV-Hypr strain and advance our understanding of mechanisms within the CNS that may be involved in vaccination breakthroughs.

## Results

### TP21 and TBEV-280 immunization protects mice from lethal TBEV-Hypr infection

First, we tested the efficacy of immunity induced by pre-immunization with TP21 or TBEV-280 to confer protection against lethal TBEV-Hypr infection. As demonstrated in Figure 1, all mock-treated mice challenged with TBEV-Hypr started showing clinical signs from 7 dpi onward with body weight loss accompanied by signs of weakness, reduced activity, pilo-erection, kyphosis, and lethargy, often combined with neurological indicators of ataxia and paresis of hind legs. Nearly 50% of mice attained clinical scores corresponding to humane endpoint at 8 dpi and the rest had to be euthanized due to severe disease by 10 dpi. Notably, all mice that were immunized with TP21 or TBEV-280 were protected from developing clinical signs and survived lethal TBEV infection until study endpoint (14 dpi) with no notable changes in the body weights. These results show that pre-immunization with TP21 or TBEV-280 is effective in inducing immunity in mice, which conferred protection from clinical and lethal TBEV infection.

**Figure 1.**
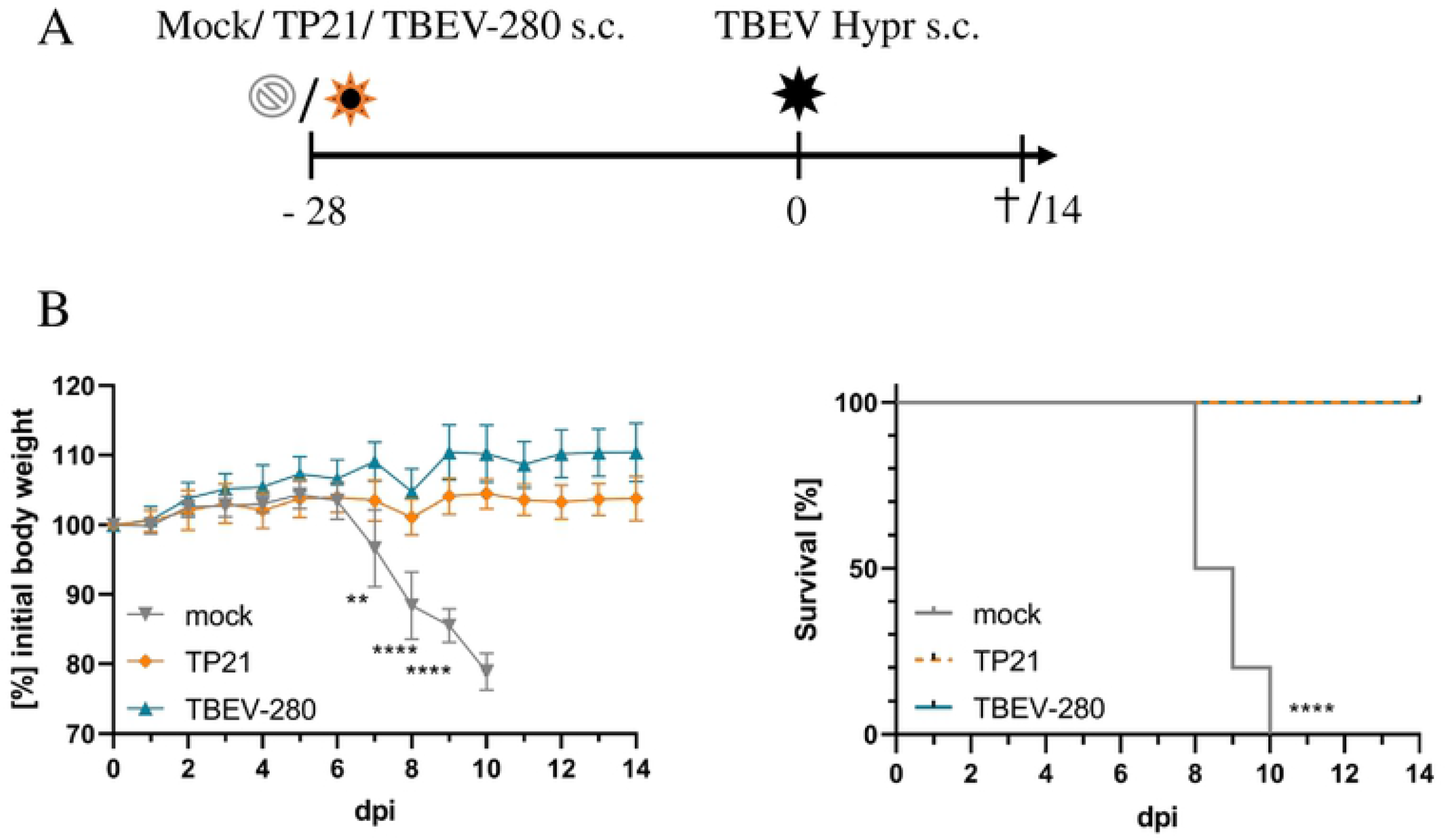
Low-virulent LGTV (TP21) and naturally avirulent TBEV (TBEV-280) immunization protects mice from highly pathogenic TBEV (TBEV-Hypr) challenge. **(A)** Schematic representation of the study. **(B)** Changes in body weight and the survival of the mice post TBEV-Hypr challenge was monitored for a period of 14 days. Data is presented as mean ± SD (n=10) of percentage change relative to initial body weight on the day of TBEV-Hypr challenge. Body weight data was analysed using unpaired students t-test and survival data analysed by log-rank test (** p< 0.01, **** p< 0.0001).

### Effect of TP21 and TBEV-280 immunization on TBEV-Hypr replication in organs

To gain insight into the virus control in immunized mice, we compared distribution of TBEV-Hypr in peripheral organs and in the CNS of the mock-treated, TP21 or TBEV-280 immunized mice after subsequently infected with TBEV-Hypr. As assessed by qPCR analysis designed to detect TBEV RNA, high copy numbers of viral RNA were found in the spleen and brain of mock-immunized mice that were sacrificed between 8-10dpi due to high clinical scores (Figure 2A). Furthermore, neuro-tropism and ability of TBEV-Hypr to predominantly replicate in neural tissue was evident from highest virus load observed in brains as compared to spleens of control mice (Figure 2A). At the study endpoint (14dpi), viral RNA was not detected in the spleens of any of the mice that experienced a prior immunization with TP21 or TBEV-280, which shows effective protection of peripheral organs from virus replication. In contrast, significant numbers of viral RNA copies were detected in the brains of some of the TP21 and all of the TBEV-280 pre-immunized mice (Figure 2A). However, viral copy numbers detected in these mice were significantly lower than those of mock control mice. Similar observations were made by qPCR analysis of the spinal cords (Supplementary Figure S1). TBEV primers used in this assay do not detect LGTV RNA at numbers less than 10^5^ copies(data not shown). Since viral copy numbers detected in LGTV immunized mice are below detection limit we can confirm that only TBEV RNA is detected in these mice. Immunostaining of brain sections for TBEV antigen confirmed presence of TBEV-Hypr infected cells across different regions of brains of mock control mice. Cerebral cortex, thalamus, hypothalamus and pons appeared to be the regions with highest viral loads (Figure 2B-C). Interestingly, no TBEV antigen was detected in TP21 or TBEV-280 immunized mouse brains (Figure 2D-E). Taken together these results confirm that TP21 and TBEV-280 immunization qPCR positive induces protective immune response against TBEV-Hypr and effectively keeps virus replication and spread under control. Apparently, virus escaping from peripheral immunity may enter the CNS and replicate there before being cleared by local inflammatory or immune mechanisms.

**Figure 2.**
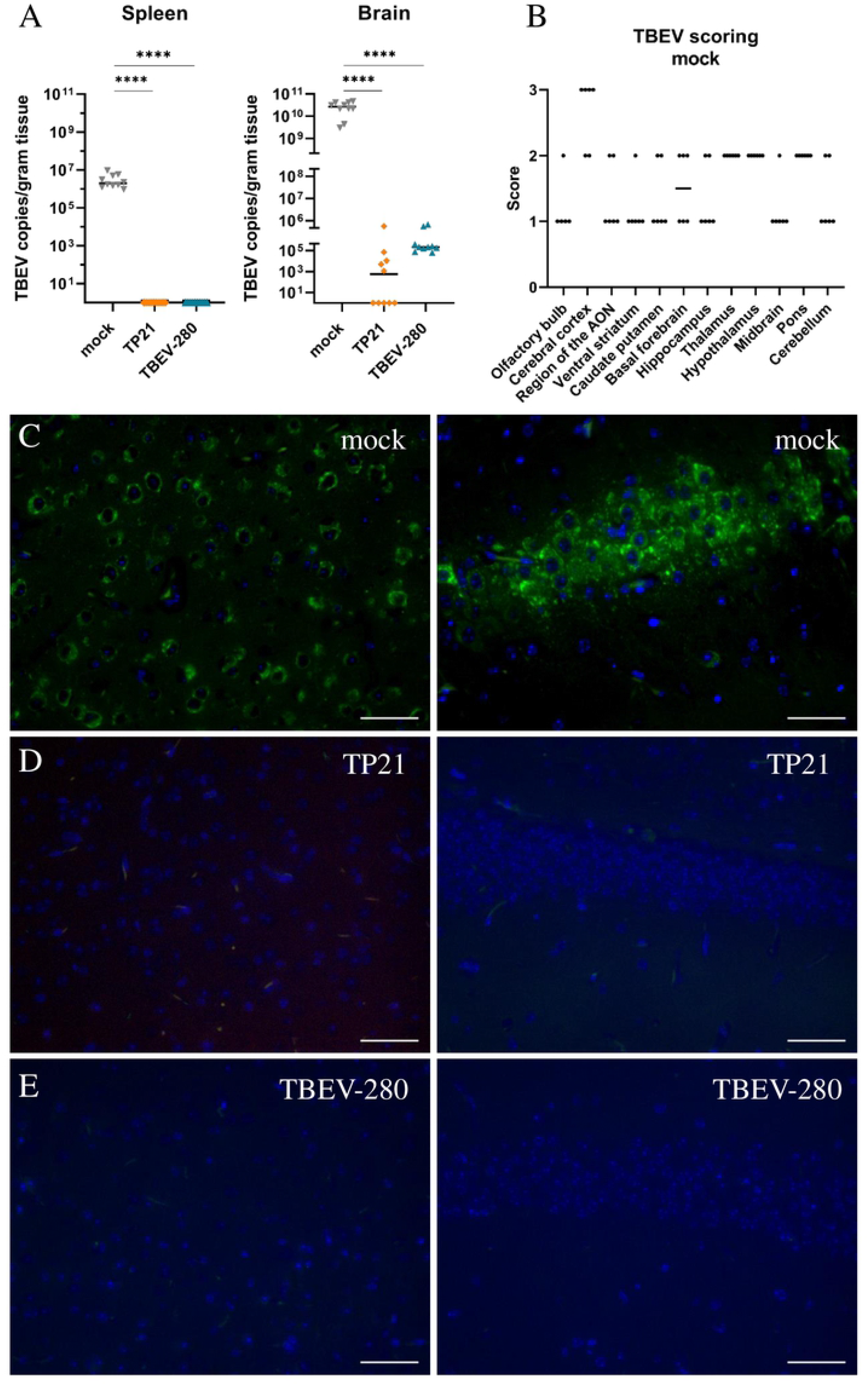
TBEV distribution in the periphery and in the CNS. (**A)** qPCR analysis of viral RNA isolated from the spleen and brain of mock, TP21 and TBEV-280 immunized mice. RNA copies per gram tissue were determined and data obtained from individual mice were plotted, bars indicate median (n=10; ****p < 0.0001) **(B)**. Tissue sections were stained with antibody targeting TBEV E-protein (green) and Hoechst (nuclei, blue) and virus antigen distribution in brain of mock immunized mice was scored based on the frequency of infected cells in the HPF analyzed (n=6). Representative immunoflourescence images from cerebral cortex (left panels) and hippocampus (right panels) regions from mock immunized **(C)** TP21 immunized **(D)** and TBEV-280 immunized **(E)** mice infected with TBEV-Hypr are depicted here. Scale bars 50µm.

### TBEV infects neurons

Immunofluorescence staining for TBEV antigen of brain sections from control, TBEV-Hypr-infected mice revealed high numbers of infected cells across different brain regions. Based on location and morphology, most of the infected cells appeared to be neurons (Figure 3A-B), as has been reported previously (22,23). However, to further investigate whether major glial cell types such as microglia and astrocytes could also constitute a target of TBEV, double labeling of brain sections using antibodies targeting TBEV antigen and microglia/macrophages (Iba-1^+^) or astrocytes (GFAP^+^) was performed. Interestingly, none of the regions that were screened displayed astrocytes (GFAP^+^ cells) that co-localized with TBEV antigen (Figure 3C-D). Largely, microglia/macrophage (Iba-1^+^cells) also failed to show co-localized TBEV antigen. However, in some areas of cerebral cortex and midbrain of single TBEV-Hypr infected control mice we detected Iba-1^+^ cells co-stained with TBEV antigen (Figure 3E-F). Collectively these results confirm that neurons constitute the primary target of TBEV infection.

**Figure 3.**
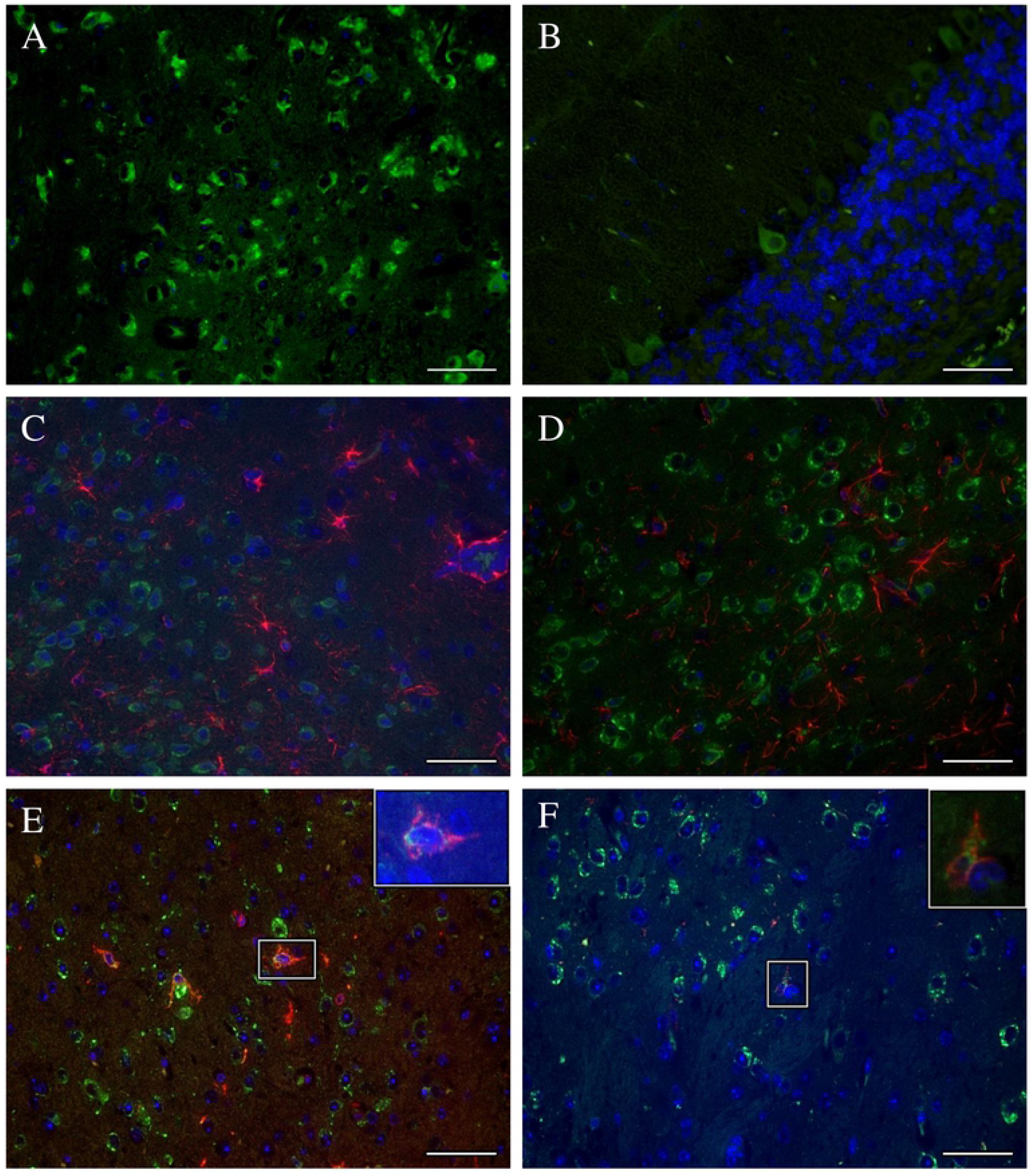
Immunoflourescence determined the cellular targets of TBEV infection in brain. Brain slices were stained using antibody targeting TBEV E-protein (green) and Hoechst (nuclei, blue). Reperesentative images from thalamus (A) and cerebellum (B) show infected cells that morphologically resemble neurons. Double labeling of TBEV E-protein (green) and glial fibrillary acid protein (GFAP, astrocytes; red) of hypothalamus (**C**) and cerebral cortex (D) shows no colocalization of TBEV antigen with astrocytes. Co-stainings with anti-Iba-1 (microglia/macrophages marker; red) and TBEV E-protein (green) show mostly uninfected microglia/macrophages. However, few Iba-1^+^ TBEV^+^ cells detected in the midbrain (E) and thalamus (F) of mock immunized and TBEV-Hypr infected mice. Scale bars 50µm.

### Marked neuronal necrosis observed after TBEV-Hypr infection in mock control mice

The results so far indicate that TBEV infects and replicate within neurons upon entering the CNS. Histopathological analysis of HE stained brain sections revealed significant neuronal necrosis in mock immunized mice that were challenged with TBEV-Hypr (Figure 4A-B). The extent of neuronal damage was assessed on a scale of 3 based on the presence of single scattered necrotic neurons per HPF (score 1) or greater than 30% affected neurons per HPF (score 3) in the regions evaluated. Neuronal damage was more prominent in the olfactory bulb, cerebral cortex and midbrain regions of heavily infected brains of mock treated mice. Interestingly, neuronal necrosis was also detected in thalamus, hypothalamus and pons of some of the TP21 immunized mice and occasionally in the cerebral cortex of TBEV-280 immunized mice (Figure 4C and Supplementary Figure S2). This indicates that the TBEV specific immune response induced by TP21 and TBEV-280 immunization protects mice from clinical signs but does not fully protect all immunized mice from neuronal damage.

**Figure 4.**
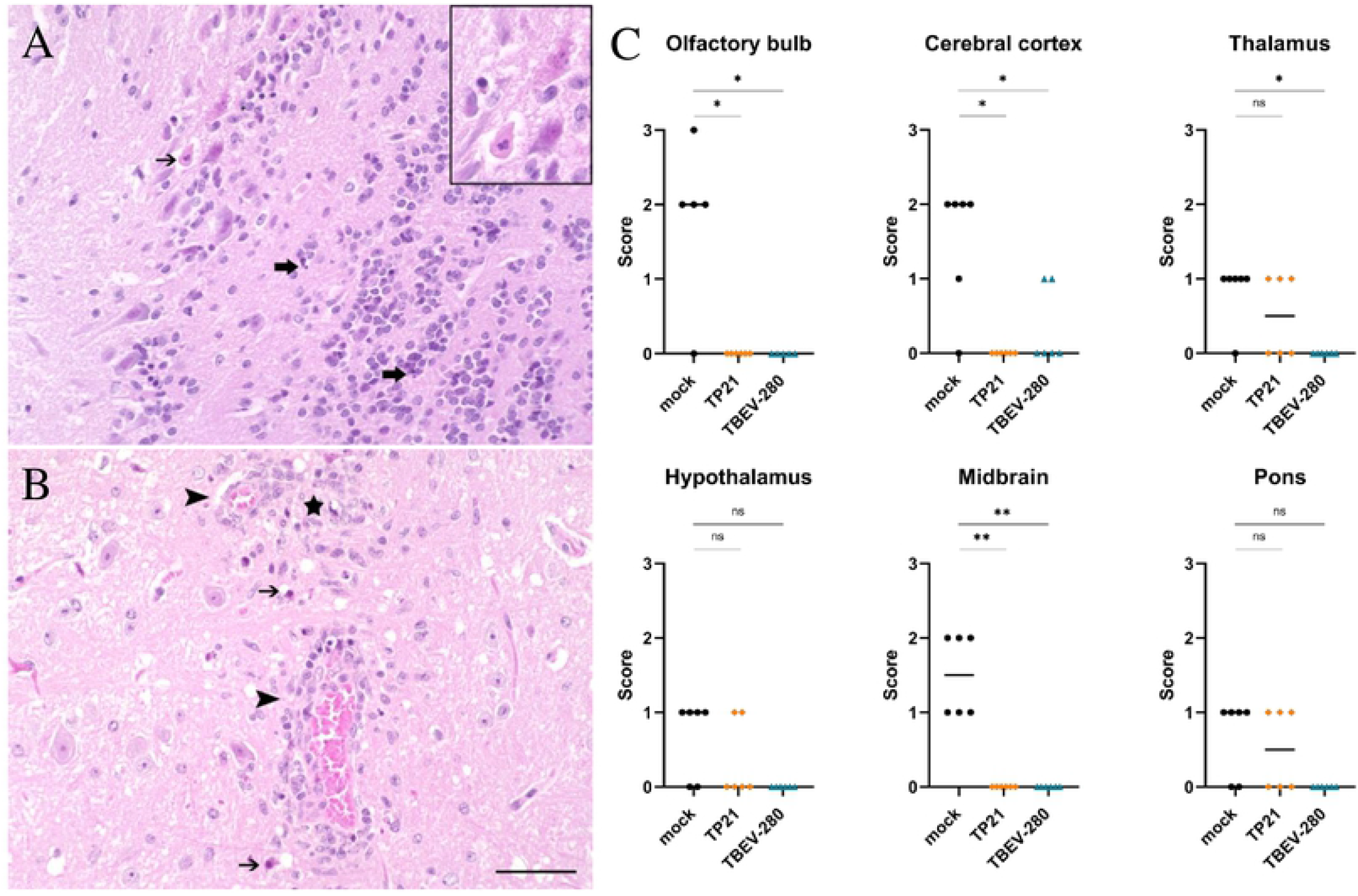
Assessment of neuronal damage in brain following TBEV-Hypr challenge. Representative images of HE staining from olfactory bulb (A) and thalamus (B) of mock immunized and TBEV-Hypr infected mice demonstrating neuronal necrosis (thin arrows), inflammation (arrow heads), cellular debris/ pyknotic nuclei (thick arrows) and gliosis (star) are shown here; Scale bars 50µm. (C) Neuronal necrosis in different regions of brain of mock, TP21 and TBEV-280 immunized and TBEV-Hypr infected mice (n=6) have been scored on a scale of 0 to 3 (see methods) and presented here (*p < 0.05, **p < 0.01).

### Inflammation and gliosis in the brain

Neuroinfection often results in extensive inflammation within the CNS which is usually characterized by perivascular infiltrates (PVI) of recruited peripheral inflammatory cells as well as hypertrophy and hyperplasia of local glial cells (gliosis). Such changes are not observed in brain of naïve mice. Examination of HE stained brain sections revealed significantly more PVI of mononuclear cells across different regions of brains of mock-immunized mice compared to TP21- and TBEV-280-immunized mice (Figure 5 & Supplementary Figure S3). Similarly, areas of gliosis were more prominent in mock immunized mice than in TP21 and TBEV-280 immunized mice (Figure 6 and Supplementary Figure S4), in line with the attenuated course of disease after immunization. Interestingly, inflammatory infiltrates as well as multifocal, mild accumulation of glial cells were also detected within the brains of TP21 and TBEV-280 immunized mice. This again shows that TBEV possibly gained access to the CNS despite the presence of peripheral protective immunity and triggers neuroinflammation.

**Figure 5.**
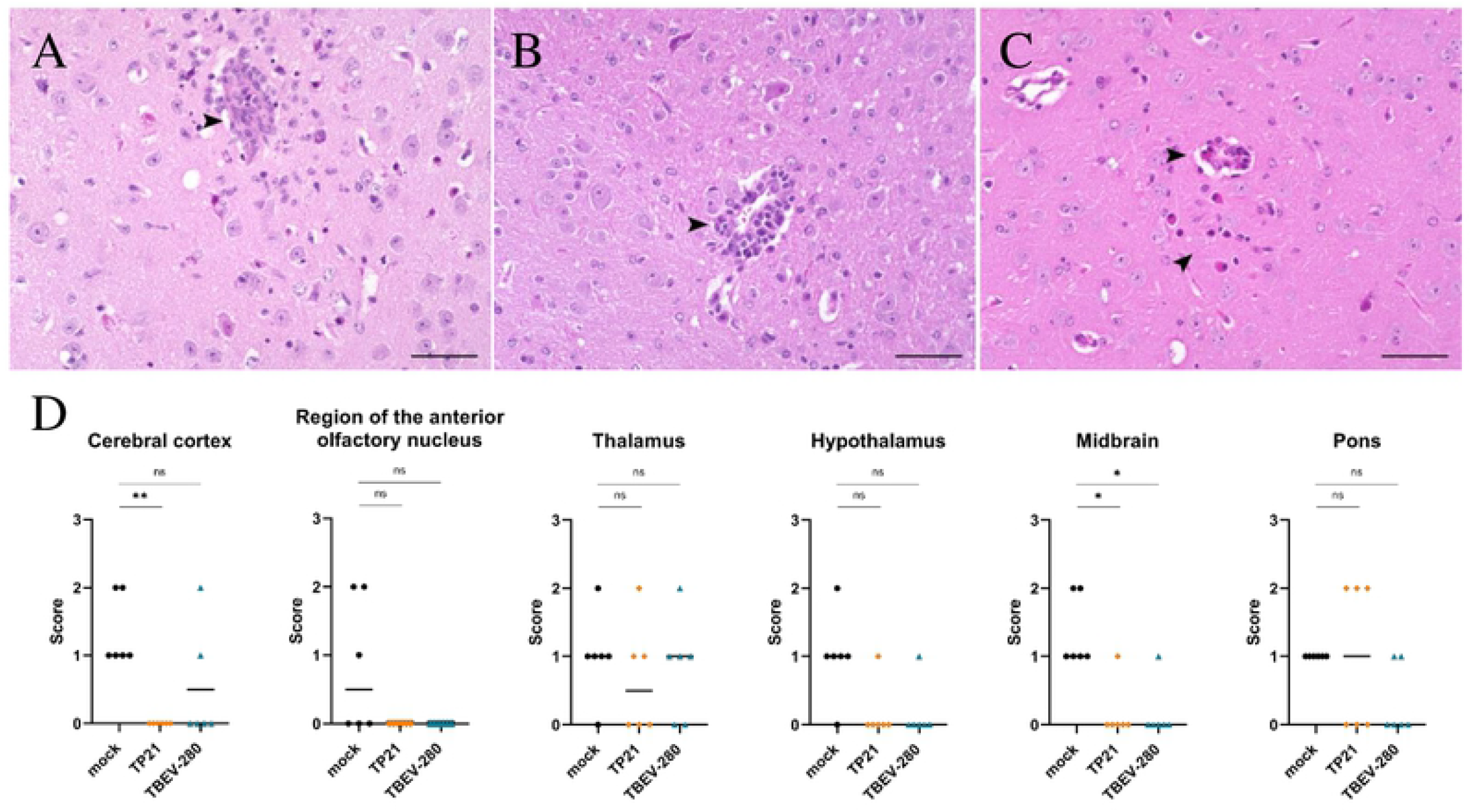
Inflammation in the brain. Regions with perivascular infiltrates (PVI) were assessed using HE staining. PVI (arrow heads) were not only detected in mock (A) but also in TP21 **(B)** and TBEV-280 **(C)** immunized mice that were subjected to TBEV-Hypr challenge; scale bars 50µm. (D). High power fields from different regions of brain of mock, TP21 and TBEV-280 immunized mice (n=6) were screened and the score (see methods) obtained for PVI from individual mice was plotted here. (*p < 0.05; **p < 0.01).

**Figure 6.**
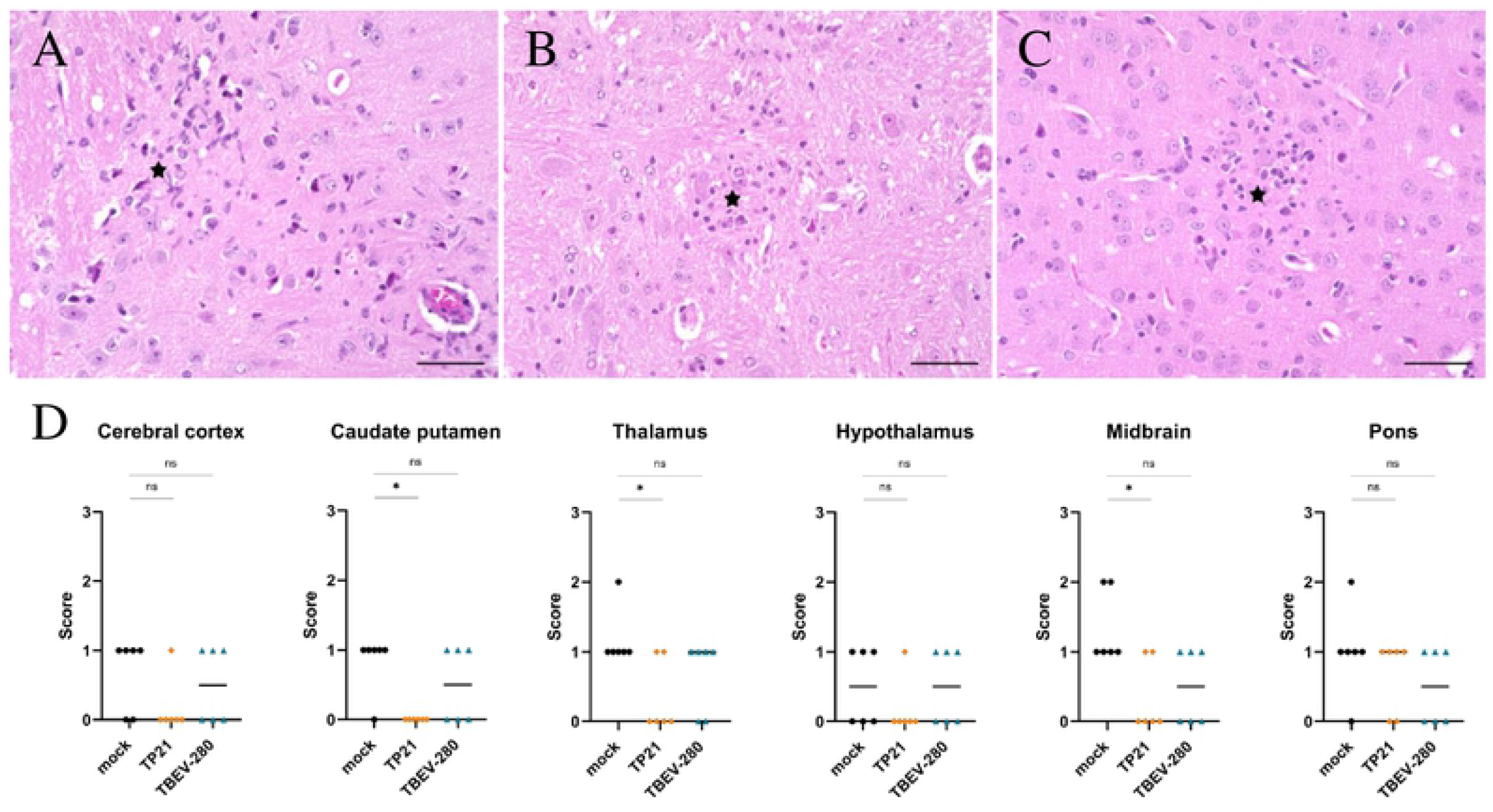
Reactive changes in the brain following TBEV-Hypr challenge. Gliosis (star) was detected in the brain of mock **(A)** TP21 **(B)** and TBEV-280 **(C)** immunized mice that were subjected to TBEV-Hypr challenge; scale bars 50µm. (D) Gliosis was scored on a scale of 0 to 3 (see methods) by screening HPF from different brain regions of mock, TP21 and TBEV-280 immunized mice (n=6; *p < 0.05).

## Discussion

In the present study, we have shown that immunization of mice with LGTV, a closely related low-virulent flavivirus, or a naturally avirulent TBEV strain (TBEV-280) provided protective immunity in mice against symptomatic and lethal infection with TBEV-Hypr strain of TBEV. Interestingly, traces of viral RNA and signs of neuroinflammation were found in the brains of LGTV and TBEV-280 immunized mice that did not display any clinical signs upon lethal TBEV-Hypr challenge. This is of particular interest in the light of breakthrough TBE that have been reported in vaccinated individuals (24,25).

Neuronal cells are regarded as primary targets of TBEV (22,23). Accordingly, in the absence of vaccine-induced protective immunity, high numbers of infected neurons were detected by immunostaining in TBEV-Hypr infected mice. As we could not detect viral antigen within neurons of immunized mice, our initial assumption was that immune response induced by LGTV and TBEV-280 pre-immunization effectively cleared TBEV-Hypr from the periphery and prevented its spread into the CNS. However, detailed qPCR analyses of brain and spinal cord from these mice reflected a different outcome. Indeed, the average viral RNA copy number detected in TP21 or TBEV-280 immunized mice were several folds lower when compared to the numbers detected in mock immunized mice. The observed viral RNA copy numbers by qPCR in the brains of the TP21 or TBEV-280 immunized mice (to the order of 10^5^ genome copies/g) were significantly higher than could be expected without active virus TBEV replication. Hence, the most likely explanation for the presence of high viral RNA copy numbers in brains and spinal cords of the immunized mice is that TBEV-Hypr is able to enter the CNS of immunized mice and probably replicate within the neural tissue. This hypothesis is furthermore supported by the presence of inflammatory (PVI) and reactive (gliosis) changes within the CNS of vaccinated mice, which both constitute frequent findings during and after neuroinfection (18). Besides neurons, some studies have demonstrated TBEV infection of astrocytes (23,26,27). Furthermore, it has been shown that TBEV can infect and survive in rat and human astrocytes for several days without affecting each other’s viability (26,27). However, co-staining of brain tissue sections did not provide evidence of TBEV antigen within astrocytes either in mock-, LGTV- or TBEV-280-immunized mice after challenge. Similarly, only isolated microglia cells in certain brain regions in mock immunized mice showed co-labeling of Iba-1 with TBEV antigen. Microglia are highly efficient phagocytes and actively scavenge cell debris in the event of tissue damage. Therefore, it remains unclear whether detected signal resulted from infection of Iba1-positive cells or from phagocytosis. There is evidence that neurotropic flaviviruses can influence the behavior of microglia and astrocytes and either assist in viral clearance or augment neuropathogenesis by producing toxic neuroinflammatory mediators (17,20,28–30). To avoid any permanent damage, inflammation within the CNS is highly regulated and is primarily mediated by microglia and astrocytes. In response to any perturbation caused within the CNS, these glial cells proliferate and migrate to the affected areas and provide neuroinflammatory mediators and neurotrophic factors to limit the damage (31–33). This phenomenon coined reactive gliosis (34) is characteristic of neural tissue damage caused by injury or infection, which could be also have occurred within the present study. In mock immunized mice after TBEV-Hypr challenge, we also observed gliosis, particularly in those areas with increased viral load. Interestingly, mild areas of gliosis have also been observed in certain brain regions of the TP21 and TBEV-280 immunized mice after TBEV-Hypr challenge, indicating reactive changes within CNS even after immunization. Since we could not detect TBEV antigen in these brains it is difficult to conclude that the areas with gliosis directly correspond to those with assumed previous virus infection. Taken together these findings are highly indicative of TBEV gaining access to the CNS and triggering inflammation despite of pre-existing TP21 or TBEV-280 induced immunity. We have observed the presence of PVI in cerebral cortex, thalamus, hypothalamus and midbrain regions of mock immunized mice. In contrast, PVIs were limited to certain brain regions in immunized mice. This again may reflect differences in virus distribution within the CNS of mock immunized and immunized mice.

Extensive neuronal necrosis was found in infected brains of mock immunized mice. There was severe loss of neurons especially in the olfactory bulb, cerebral cortex and midbrain regions accompanied by detection of numerous neurons staining positive for TBEV antigen by IF as well as inflammatory and reactive changes. However, it is not clear if the observed neuronal damage represents a direct effect of virus infection or a secondary consequence of inflammation. It has been shown for other flaviviruses that infection and virus replication can trigger apoptosis and necrosis in neurons (19,35–40). Similarly, inflammation as a consequence of neuroinfection can also caus substantial neuronal loss (41,42).

Probably the most intriguing finding from this study is the detection of neuronal necrosis combined with inflammation and gliosis despite of vaccine-induced protective immunity to TBEV in some mice. This phenomenon was more prominent in TP21 than in TBEV-280 immunized mice. This is of special interest, considering breakthrough TBE in vaccinated individuals. The observation that TBEV may still enter the CNS despite vaccine induced protective immunity while causing inflammation and neuronal damage may at least in part explain vaccination breakthroughs in TBE vaccinated individuals.

In conclusion, this study demonstrates for the first time the presence of TBEV in the CNS of immunized mice that are protected from lethal or clinically manifest infection. However, absence of detectable TBEV antigen in these brains via IF hints at an important role of local inflammatory mediators in checking the virus spread within the CNS in the event virus escapes host immune response in the periphery. Nonetheless, the actual cause of the neuronal necrosis in immunized mice remains unclear. Whether this is caused by virus infection directly or a consequence of subsequent inflammation or even the action of infiltrating virus-specific T cells remains a matter of further investigation. Recently Garber *et al*. demonstrated that ZIKV specific T cells through microglia mediate neuronal loss and hence result in cognitive dysfunction (17). Furthermore, CD8^+^ T cells that clear infected cells also cause neuronal damage (18,19,43). Finally, it is tempting to speculate whether subtle neuronal necrosis as observed in immunized mice is the reflection of a broader challenge of raising sufficient immune mediated protection of the CNS from TBEV invasion, which may also be the basis of current human vaccination failures.

## Material and Methods

### Mice and ethics

Six-week old female C57BL/6JOlaHsd (BL6) mice were obtained from Envigo, Inc. (Indiana, USA). Mice were housed in isocage systems with individually ventilated cages. Experiments were done in biosafety level 3 laboratories of the Institute of Parasitology, Biology Center of Czech Academy of Sciences, České Budějovice, Czech Republic. The protocol was approved by the Departmental Expert Committee for the Approval of Projects of Experiments on Animals of the Czech Academy of Sciences and the Committee on the Ethics of Animal Experimentation at the Institute of Parasitology (permit No. 29/2016). All experiments were done in accordance with Czech national law guidelines (animal Welfare Act No. 246/1992 Col.) and European Union guidelines for work with animals.

### Viruses

LGTV strain TP21 (referred to as TP21) was isolated 1956 from a pool of hard ticks (*Ixodes granulatus*) of forest rats near Kuala Lumpur, Malaysia (44). TP21 was grown in Vero B4 and Vero E6 cells and titers were determined by TCID50 assay on Vero E6 cells. Virus was provided the Department of Molecular Immunology, Helmholz Centre for Infection Research, Braunschweig, Germany. Highly virulent TBEV strain TBEV-Hypr (further on described as TBEV-Hypr) was isolated in 1953 from a diseased child in Brno (former Czechoslovakia). The virus was passaged eight times in suckling mice brains before its use in this study. A naturally avirulent strain TBEV-280 was isolated in 1987 from a pool of *I. ricinus* ticks near Kaplice (former Czechoslovakia). The virus was plaque-purified three times, and passaged three times in suckling mice brains, one time in UFK-NB4 cells and one times in SK-N-SH cells before its use in this study. TBEV titers were determined by plaque assay as described in (45). The TBEV strains were provided by the Collection of Arboviruses, Institute of Parasitology, Biology Centre of the Czech Academy of Sciences, České Budějovice, Czech Republic.

### Immunization study

Subcutaneous (s.c) administration of 500 pfu of TBEV-Hypr results in 100% lethality in mice. Here, we utilized this mouse model to test the efficacy of TP21 and TBEV-280 in providing protection against lethal TBEV-Hypr challenge. At 28 days prior to TBEV-Hypr challenge, mice were administered s.c with medium (mock), TP21 (10^4^ pfu) or TBEV-280 (10^4^ pfu). Following a s.c. challenge with 500 pfu of TBEV-Hypr, development of clinical signs in mice was monitored for a period of 14 days and when clinical score corresponding to humane endpoint was reached, they were sacrificed.

### RNA isolation and quantitative real-time PCR (qPCR)

To determine viral loads in organs, half the brain, spinal cord and spleen were dissected, weighted and homogenized in AVL Buffer (Qiagen). RNA was isolated using QIAamp viral RNA Mini QC Kit (Qiagen) and the QIAcube machine. Viral load was quantified by Taq-Man qPCR using OneStep RT-PCR Kit (Qiagen) and AriaMx Real-time PCR Systems (Agilent). TBEV forward primer (5’-3’ GGGCGGTTCTTGTTCTCC), TBEV reverse primer (5’-3’ ACACATCACCTCCTTGTCAGACT) and TBEV probe (5’-3’ TGAGCCACCATCACCCAGACACA) were designed by (46). AriaMX software (version 1.5; Agilent) in combination with intraassay TBEV standard curve was used to analyse the data and viral copies were extrapolated by individual organ weight in RNA copies/gram.

### Histology

Brains were cut midsagittal and fixed for 48 hours in 4% Histofix (Carl Roth). After fixation organs were stored in PBS till paraffin embedment and cut of serial sections on a microtome (2-3 µm; Leica RM 2035; Leica Instruments GmbH, Nuβloch, Germany). Sections were stained by hematoxylin and eosin (HE) and immunofluorescence (IF).

### Immunofluorescence

IF was performed as described before (47). Briefly, sections of formalin-fixed, paraffin-embedded brains were dewaxed and rehydrated using graded alcohols. For antigen retrieval, sections were heated in a microwave oven (800 W) for 20 min in 0.01 M citrate buffer, followed by application of inactivated goat serum or horse serum, respectively. Afterwards, sections were incubated with primary antibodies (see Table 1) targeting TBEV E-protein (clone 1493) (48), glial fibrillary acidic protein (GFAP, astrocytes) and ionized calcium-binding adapter molecule (Iba-1, microglia/macrophages) overnight at 4°C. For detecting TBEV-infected astrocytes and microglia, double labeling of TBEV E-protein with GFAP and Iba-1 was performed. Negative control sections were incubated with equally diluted rabbit or mouse serum. For visualization of antigen-antibody reactions, sections were treated with Alexa Fluor 488- or Cy3-labeled secondary antibodies (1:200) for 1 hour at room temperature. Nuclei were stained with bisbenzimide (Hoechst 33258, 0.01 % in methanol, 1:100; Sigma-Aldrich, Taufkirchen, Germany) and mounted with fluorescence mounting medium (Dako Diagnostika, Hamburg, Germany). Sections were evaluated with a fluorescence microscope (Keyence BZ-9000E, Keyence Deutschland GmbH, Neu-Isenburg, Germany).

**Table 1.**
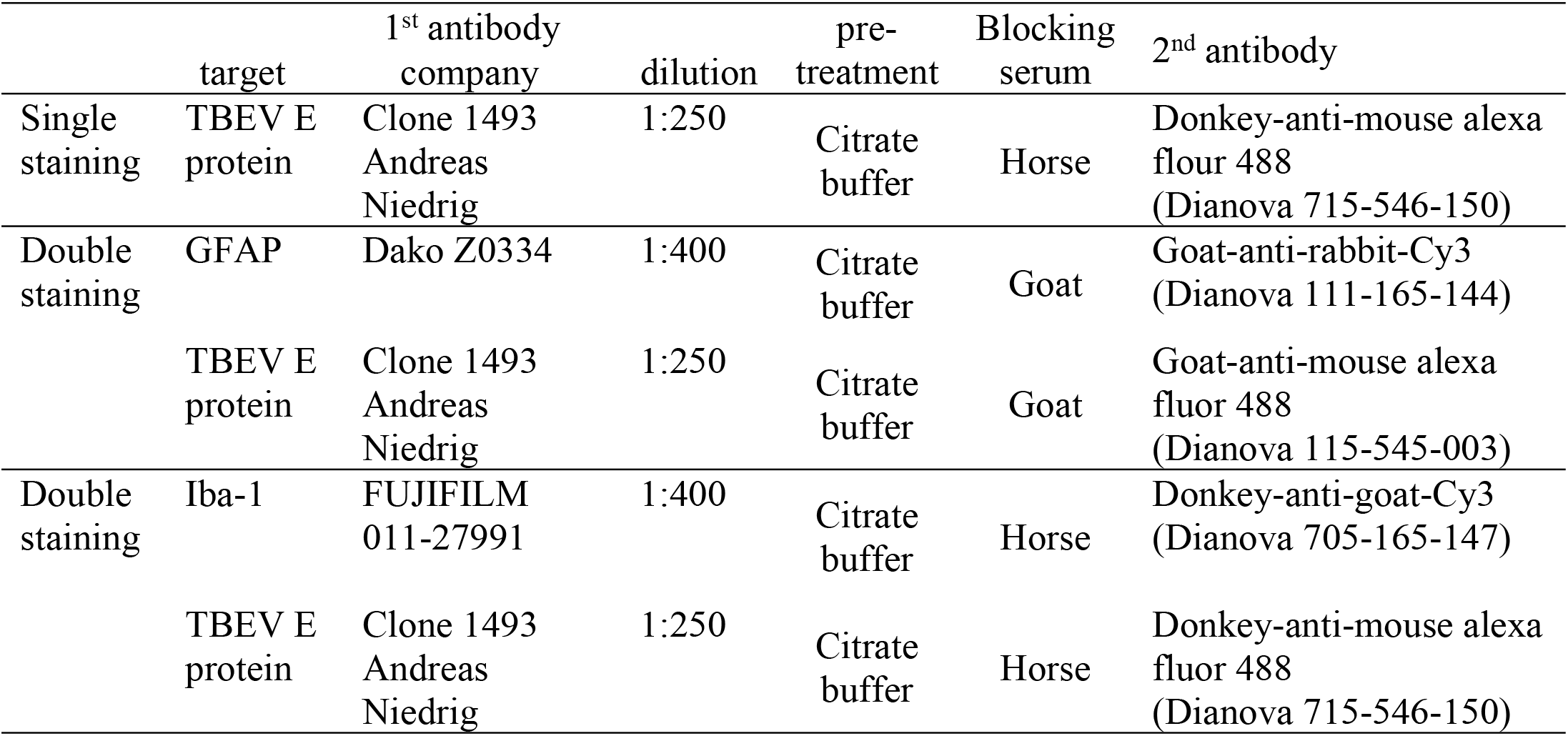
Antibodies and reagents of immunofluorescence

|  | target | 1<sup>st</sup> antibody<br>company | dilution | pre-<br>treatment | Blocking<br>serum | 2<sup>nd</sup> antibody |
| --- | --- | --- | --- | --- | --- | --- |
| Single<br>staining | TBEV E<br>protein | Clone 1493<br>Andreas<br>Niedrig | 1:250 | Citrate<br>buffer | Horse | Donkey-anti-mouse alexa<br>fluor 488<br>(Dianova 715-546-150) |
| Double<br>staining | GFAP | Dako Z0334 | 1:400 | Citrate<br>buffer | Goat | Goat-anti-rabbit-Cy3<br>(Dianova 111-165-144) |
|  | TBEV E<br>protein | Clone 1493<br>Andreas<br>Niedrig | 1:250 | Citrate<br>buffer | Goat | Goat-anti-mouse alexa<br>fluor 488<br>(Dianova 115-545-003) |
| Double<br>staining | Iba-1 | FUJIFILM<br>011-27991 | 1:400 | Citrate<br>buffer | Horse | Donkey-anti-goat-Cy3<br>(Dianova 705-165-147) |
|  | TBEV E<br>protein | Clone 1493<br>Andreas<br>Niedrig | 1:250 | Citrate<br>buffer | Horse | Donkey-anti-mouse alexa<br>fluor 488<br>(Dianova 715-546-150) |

### Histological evaluation

HE stained slides were evaluated by semi-quantitative scoring of neuronal necrosis, gliosis and inflammation for individual brain regions. The applied scoring system included three categories. Neuronal necrosis scores ranged from 0 to 3 (0 normal; 1 single necrotic neurons, 2 less than 30 % affected neurons; 3 more than 30 % affected neurons). Gliosis, interpreted as hyperplasia and hypertrophy of glial cells, ranged from 0 to 3 (0 normal; 1 multifocal, mild; 2 multifocal, moderate; 3 multifocal, severe) and inflammation scores ranged from 0 to 3 (0 normal; 1 single perivascular infiltrates; 2 2-3 layers of perivascular infiltrates; 3 more than 3 layers of perivascular infiltrates). TBEV scoring ranged from 0 to 3 (1 <30 % positive cells, 2 30-60 % positive cells; 3 >60 % positive cells). All scorings were performed using a defined area of high power fields (HPFs; 400x).

### Statistics

Data was analyzed by GraphPad Prism Software 9. Survival curves are displayed by Kaplan-Meier curves using log rank test. Body weights are expressed by mean ± standard error (SD) in graphs and were analyzed using unpaired, two-tailed student’s - t test. Differences between viral load or HE scorings were analyzed using Mann-Whitney *U* test unless described differently. A p-value of < 0.05 was considered significant and indicated by (*).

## Acknowledgements

We thank Prof. Matthias Niedrig former employee of the Department of Virology of the Robert-Koch-Institute Berlin, Germany for providing the antibody against TBEV for these studies. We thank the Federal Ministry of Education and Research for funding this study within the TBENAGER grant for AO. This work was funded in part by the Alexander von Humboldt Foundation in the framework of the Alexander von Humboldt Professorship endowed by the German Federal Ministry of Education and Research. We thank the Deutsche Forschungsgemeinschaft (DFG, German Research Foundation) – 398066876/GRK 2485/1, VIPER for funding parts of the study. We thank for a support from the Czech Science Foundation (grant No. 20-14325S to DR, and No. 20-30500S to MP).

## Supplementary information

***Supplementary Figure S1***. Taq-Man qPCR analysis of the spinal cord of mock, TP21 and TBEV-280 immunized mice that were infected with TBEV-Hypr.

***Supplementary Figure S2***. Evaluation of neuronal necrosis by HE staining in other brain regions. Regions were scored from 0 to 3 (see methods) and statistical analyzed by Mann-Whitney test (*p < 0.05; n=6/group).

***Supplementary Figure S3***. Evaluation of inflammation using HE staining in different brain regions. Regions were scored from 0 to 3 (see methods) and statistical analyzed by Mann-Whitney test (*p < 0.05; n=6/group).

***Supplementary Figure S4***. Evaluation of gliosis using HE staining in different brain regions. Regions were scored from 0 to 3 (see methods) and statistical analyzed by Mann-Whitney test (*p < 0.05; n=6/group).

## Abbreviations

CNS: Central nervous system
dpi: Days post infection
EMEM: Eagle’s Minimum Essential medium
GFAP: Glial fibrillary acidic protein marker
HE: Haematoxylin and eosin staining
HPF: High power fields
Iba-1: Ionized calcium binding adaptor molecule 1 marker
IHC: Immunohistochemistry
LAV: Live-attenuated vaccine
LGTV: Langat virus
pfu: Plaque-forming unit
PVI: Perivascular infiltrates
qPCR: Quantitative real-time PCR
s.c.: Subcutaneous
TBE: Tick-borne encephalitis
TBEV: Tick-borne encephalitis virus
ZIKV: Zika virus

## References

1. Mandl CW, Ecker M, Holzmann H, Kunz C, Heinz FX. Infectious cDNA clones of tick-borne encephalitis virus European subtype prototypic strain Neudoerfl and high virulence strain Hypr. J Gen Virol. 1997;78(5):1049–57.

2. Gritsun TS, Lashkevich VA, Gould EA. Tick-borne encephalitis. Antiviral Res [Internet]. 2003;57(1–2):129–46.

3. Dai X, Shang G, Lu S, Yang J, Xu J. A new subtype of eastern tick-borne encephalitis virus discovered in Qinghai-Tibet Plateau, China. Emerg Microbes Infect. 2018;7(1):74.

4. Kovalev SY, Mukhacheva TA. Reconsidering the classification of tick-borne encephalitis virus within the Siberian subtype gives new insights into its evolutionary history. Infect Genet Evol [Internet]. 2017;55:159–65.

5. Blom K, Cuapio A, Sandberg JT, Varnaite R, Michaelsson J, Bjorkstrom NK, et al. Cell-Mediated Immune Responses and Immunopathogenesis of Human Tick-Borne Encephalitis Virus-Infection. Front Immunol. 2018;9:2174.

6. Růžek D, Dobler G, Mantke OD. Tick-borne encephalitis: Pathogenesis and clinical implications. Travel Med Infect Dis [Internet]. 2010;8(4):223–32.

7. Heinz FX, Holzmann H, Essl A, Kundi M. Field effectiveness of vaccination against tick-borne encephalitis. Vaccine. 2007;25(43):7559–67.

8. Kubinski M, Beicht J, Gerlach T, Volz A, Sutter G, Rimmelzwaan GF. Tick-borne encephalitis virus: A quest for better vaccines against a virus on the rise. Vaccines. 2020;8(3):1–45.

9. Beran J, Lattanzi M, Xie F, Moraschini L, Galgani I. Second five-year follow-up after a booster vaccination against tick-borne encephalitis following different primary vaccination schedules demonstrates at least 10 years antibody persistence. Vaccine. 2019;37(32):4623–9.

10. Minor PD. Live attenuated vaccines: Historical successes and current challenges. Virology [Internet]. 2015;479–480:379–92.

11. Seligman SJ, Gould EA. Live flavivirus vaccines: Reasons for caution. Lancet [Internet]. 2004;363(9426):2073–5.

12. Rumyantsev AA, Murphy BR, Pletnev AG. A tick-borne Langat virus mutant that is temperature sensitive and host range restricted in neuroblastoma cells and lacks neuroinvasiveness for immunodeficient mice. J Virol. 2006;80(3):1427–39.

13. Mitrova E, Mayer V. A live vaccine against tick-borne encephalitis; integrated studies, H. Histopathology of mice peripherally immunized with E5 “14” virus and challenged with virulent virus. Acta Virol [Internet]. 1975;3:219–28.

14. Mayer V, Pogady J, Starek M, Hrbka J. A live vaccine against tick borne encephalitis: integrated studies. III. Response of man to a single dose of the E5’14’ clone (Langat virus). Acta Virol. 1975;19(3):229–36.

15. Gritsun TS, Frolova T V., Pogodina V V., Lashkevich VA, Venugopal K, Gould EA. Nucleotide and deduced amino acid sequence of the envelope gene of the Vasilchenko strain of TBE virus; comparison with other flaviviruses. Virus Res [Internet]. 1993;27(2):201–9.

16. Turtle L, Bali T, Buxton G, Chib S, Chan S, Soni M, et al. Human T cell responses to Japanese encephalitis virus in health and disease. J Exp Med. 2016;213(7):1331–52.

17. Garber C, Soung A, Vollmer LL, Kanmogne M, Last A, Brown J, et al. T cells promote microglia-mediated synaptic elimination and cognitive dysfunction during recovery from neuropathogenic flaviviruses. Nat Neurosci. 2019;22(8):1276–88.

18. Růžek D, Salát J, Palus M, Gritsun TS, Gould EA, Dyková I, et al. CD8+ T-cells mediate immunopathology in tick-borne encephalitis. Virology [Internet]. 2009;384(1):1–6.

19. Gelpi E, Preusser M, Laggner U, Garzuly F, Holzmann H, Heinz FX, et al. Inflammatory response in human tick-borne encephalitis: analysis of postmortem brain tissue. J Neurovirol [Internet]. 2006;12(4):322–7.

20. Cornelius ADA, Hosseini S, Schreier S, Fritzsch D, Weichert L, Michaelsen-Preusse K, et al. Langat virus infection affects hippocampal neuron morphology and function in mice without disease signs. J Neuroinflammation. 2020;17(1):278.

21. Růžek D, Gritsun TS, Forrester NL, Gould EA, Kopecký J, Golovchenko M, et al. Mutations in the NS2B and NS3 genes affect mouse neuroinvasiveness of a Western European field strain of tick-borne encephalitis virus. Virology [Internet]. 2008;374(2):249–55.

22. Růžek D, Vancová M, Tesařová M, Ahantarig A, Kopecký J, Grubhoffer L. Morphological changes in human neural cells following tick-borne encephalitis virus infection. J Gen Virol [Internet]. 2009;90(7):1649–58.

23. Fares M, Cochet-Bernoin M, Gonzalez G, Montero-Menei CN, Blanchet O, Benchoua A, et al. Pathological modeling of TBEV infection reveals differential innate immune responses in human neurons and astrocytes that correlate with their susceptibility to infection. J Neuroinflammation. 2020;17:76.

24. Sendi P, Hirzel C, Pfister S, Ackermann-Gäumann R, Grandgirard D, Hewer E, et al. Fatal outcome of European tick-borne encephalitis after vaccine failure. Front Neurol. 2017;8:119.

25. Andersson CR, Vene S, Insulander M, Lindquist L, Lundkvist Å, Günther G. Vaccine failures after active immunisation against tick-borne encephalitis. Vaccine. 2010;28(16):2827–31.

26. Potokar M, Korva M, Jorgačevski J, Avšič-Županc T, Zorec R. Tick-Borne Encephalitis Virus Infects Rat Astrocytes but Does Not Affect Their Viability. PLoS One. 2014;9(1):e86219.

27. Palus M, Bílý T, Elsterová J, Langhansová H, Salát J, Vancová M, et al. Infection and injury of human astrocytes by tick-borne encephalitis virus. J Gen Virol. 2014;95(11):2411–26.

28. Myint KSA, Kipar A, Jarman RG, Gibbons R V., Perng GC, Flanagan B, et al. Neuropathogenesis of Japanese Encephalitis in a Primate Model. PLoS Negl Trop Dis. 2014;8(8):e2980.

29. Ho C-Y, Ames HM, Tipton A, Vezina G, Liu JS, Scafidi J, et al. Differential neuronal susceptibility and apoptosis in congenital Zika virus infection. Ann Neurol. 2017;82(1):121–7.

30. Pokorna Formanova P, Palus M, Salat J, Hönig V, Stefanik M, Svoboda P, et al. Changes in cytokine and chemokine profiles in mouse serum and brain, and in human neural cells, upon tick-borne encephalitis virus infection. J Neuroinflammation. 2019;16(1).

31. Prajeeth CK, Kronisch J, Khorooshi R, Knier B, Toft-Hansen H, Gudi V, et al. Effectors of Th1 and Th17 cells act on astrocytes and augment their neuroinflammatory properties. J Neuroinflammation. 2017;14(1):204.

32. Detje CN, Lienenklaus S, Chhatbar C, Spanier J, Prajeeth CK, Soldner C, et al. Upon Intranasal Vesicular Stomatitis Virus Infection, Astrocytes in the Olfactory Bulb Are Important Interferon Beta Producers That Protect from Lethal Encephalitis. J Virol. 2015;89(5):2731–8.

33. Weber E, Finsterbusch K, Lindquist R, Nair S, Lienenklaus S, Gekara NO, et al. Type I Interferon Protects Mice from Fatal Neurotropic Infection with Langat Virus by Systemic and Local Antiviral Responses. J Virol. 2014;88(21):12202–12.

34. Burda JE, Sofroniew M V. Reactive gliosis and the multicellular response to CNS damage and disease. Neuron. 2014;81(2):229–48.

35. Wang Q, Xin X, Wang T, Wan J, Ou Y, Yang Z, et al. Japanese Encephalitis Virus Induces Apoptosis and Encephalitis by Activating the PERK Pathway. J Virol. 2019;93(17).

36. Chen Z, Wang X, Ashraf U, Zheng B, Ye J, Zhou D, et al. Activation of neuronal N-methyl-d-aspartate receptor plays a pivotal role in Japanese encephalitis virus-induced neuronal cell damage. J Neuroinflammation. 2018;15(1):238.

37. Leyssen P, Paeshuyse J, Charlier N, Van Lommel A, Drosten C, De Clercq E, et al. Impact of direct virus-induced neuronal dysfunction and immunological damage on the progression of flavivirus (Modoc) encephalitis in a murine model. J Neurovirol [Internet]. 2003;9(1):69–78.

38. Růžek D, Vancová M, Tesařová M, Ahantarig A, Kopecký J, Grubhoffer L. Morphological changes in human neural cells following tick-borne encephalitis virus infection. J Gen Virol. 2009;90(7):1649–58.

39. Maximova OA, Faucette LJ, Ward JM, Murphy BR, Pletnev AG. Cellular inflammatory response to flaviviruses in the central nervous system of a primate host. J Histochem Cytochem. 2009;57(10):973–89.

40. Prikhod’ko GG, Prikhod’ko EA, Cohen JI, Pletnev AG. Infection with Langat flavivirus or expression of the envelope protein induces apoptotic cell death. Virology [Internet]. 2001;286(2):328–35.

41. Ghoshal A, Das S, Ghosh S, Mishra MK, Sharma V, Koli P, et al. Proinflammatory mediators released by activated microglia induces neuronal death in Japanese encephalitis. Glia. 2007;55(5):483–96.

42. Chen CJ, Ou YC, Lin SY, Raung SL, Liao SL, Lai CY, et al. Glial activation involvement in neuronal death by Japanese encephalitis virus infection. J Gen Virol. 2010;91(4):1028–37.

43. Shrestha B, Samuel MA, Diamond MS. CD8+ T Cells Require Perforin To Clear West Nile Virus from Infected Neurons. J Virol. 2006;80(1):119–29.

44. Gordon Smith CE. A virus resembling Russian spring-summer encephalitis virus from an ixodid tick in Malaya. Nature [Internet]. 1956;178(4533):581–2.

45. Palus M, Vojtíšková J, Salát J, Kopecký J, Grubhoffer L, Lipoldová M, et al. Mice with different susceptibility to tick-borne encephalitis virus infection show selective neutralizing antibody response and inflammatory reaction in the central nervous system. J Neuroinflammation [Internet]. 2013;10:1–13.

46. Schwaiger M, Cassinotti P. Development of a quantitative real-time RT-PCR assay with internal control for the laboratory detection of tick borne encephalitis virus (TBEV) RNA. J Clin Virol [Internet]. 2003;27(2):136–45.

47. Attig F, Spitzbarth I, Kalkuhl A, Deschl U, Puff C, Baumgärtner W, et al. Reactive oxygen species are key mediators of demyelination in canine distemper leukoencephalitis but not in theiler’s murine encephalomyelitis. Int J Mol Sci. 2019;20(13).

48. Niedrig M, Klockmann U, Lang W, Roeder J, Burk S, Dmodrow S, et al. Monoclonal antibodies directed against tick-borne encephalitis virus with neutralizing activity in vivo. Acta Virol [Internet]. 1994;38:141–9.

